# Genomic characterization and enhanced fermentation of the endophyte *Stemphylium* sp. (Aa22), a producer of bioactive alkyl-resorcinols

**DOI:** 10.1101/2025.04.02.646934

**Authors:** Jorge Rojas Lopez-Menchero, Juan Imperial, M. Fe Andrés, Carmen Elisa Díaz, Azucena González-Coloma

## Abstract

The genome of the previously described endophytic fungus *Stemphylium* sp. (strain Aa22) has been sequenced to near completion. Phylogenomic analysis placed strain Aa22 in close proximity to *Stemphylium lycopersici*. Strain Aa22 had been previously reported as the producer of the bioactive alkyl-resorcinol stemphol and derivative stempholones A and B in solid culture on rice. Genome mining for biosynthetic gene clusters (BGCs) identified 42 genomic regions predicted to encode secondary metabolites production. Among them, a single type III polyketide synthase (T3PKS)-encoding ORF (FUN_008199) was identified that shared similarity with other fungal T3PKSs. T3PKSs are responsible for the biosynthesis of alkyl-resorcinols from fatty acyl-CoA substrates. This makes the T3PKS gene a likely candidate for stempholone biosynthesis and a target for future manipulation to enhance production of bioactive alkyl-resorcinols. We also studied the production of these compounds in solid rice media and in liquid PDB medium with or without the addition of talcum powder. The highest extract yield was obtained with PDB cultures, and GC-MS analysis revealed the presence of high levels of the bioactive compound stempholone A, along with two unknown compounds (retention times of 20.96 and 24.37 min). Addition of talcum powder suppressed stempholone A production and reduced chemical diversity, with accumulation of oleamide. In contrast, the rice solid media fermentation resulted in methylated fatty acids and oleamide, with no detectable stempholone.

## Introduction

Currently, agriculture is facing the global challenge of being productive, efficient, sustainable, and environmentally friendly (1). Food production is affected by plant diseases and insect pests (2) To date, plant protection has depended on synthetic pesticides, resulting in negative effects. In this context, there has been an increase in the research on new safer bio-pesticides (3,4). Additionally, stricter pesticide safety regulations make the search for new bio-pesticides a priority (3).

Endophytic fungi are present in tissues or organs of plants without causing apparent damage (5) and contribute to the adaptation of the plant to abiotic and biotic stresses (6). Endophytic fungi are capable of producing a diverse range of bioactive compounds, including those with insecticidal, antioxidant, antifungal, antiviral, antibacterial, or cytotoxic activities that have potential as biopesticides (7–9). Thus, endophytic fungi represent a promising novel source for the biotechnological production of valuable active compounds (10–12).

Most fungal endophytes are traditionally identified morphologically and genetically through analysis of the internal transcribed spacer regions of nuclear ribosomal DNA (ITS1-5.8S-ITS4 rDNA). However, with the advent of powerful, affordable next-generation sequencing (NGS) methods, these are being increasingly used to study endophytes for taxonomic identification along with approaches such as metabolomics, metagenomics, proteomics and bioinformatics (13). Functional genomics sequencing of fungal endophytes involves a comprehensive analysis of their genetic material to understand the roles of specific genes and metabolic pathways in plant-microbe interactions. Therefore, functional genomics can reveal the potential of fungal endophytes in biotechnological applications (14).

The study of the endophytic biodiversity from medicinal plants has resulted in the identification of bioactive compounds. One example is wormwood (*Artemisia absinthium*), where endophytic bacteria are a remarkable source of antimicrobial and anticancer compounds such as 2-aminoacetophenone, 1,2-apyrazine-1,4-dione, phenazine and 2-phenyl-4-cyanopyridine (15). Furthermore, in a recent study, our group showed that the fungal endophytic strain Aa22, isolated from *A. absinthium* and previously identified as *Stemphylium solani,* was able to yield an ethyl acetate extract from the solid rice fermentation that had antifeedant properties against *Myzus persicae* aphids (16). The bioguided fractionation of this extract led to the isolation of the alkyl-resorcinols stempholone A, stempholone B and stemphol. All these compounds were aphid antifeedants, and stempholone A was also moderately nematicidal. These results highlight the potential of the endophytic strain Aa22 as a biotechnological source of natural product-based biopesticides.

Fungal fermentation conditions significantly influence the production of bioactive compounds of interest. Submerged fermentation (SmF) and solid-state fermentation (SSF) are the most widely used methods, both with advantages and disadvantages (17–19). SSF has the potential to reduce costs and energy consumption by utilizing agro-industrial waste (20). On the other hand, SmF has benefits such as higher productivity and yields (21,22), and allows for easier and more reproducible control and measurement of fermentation parameters (23). However, in SSF, the development of mycelial clumps or pellets reduces the effective transfer of oxygen and nutrient resources in the liquid phase (24). In this context, semi-solid-state fermentation (Semi-SSF) is a hybrid method that utilizes both an inert solid support and a smaller volume of culture medium (25). Semi-SSF allows the production of secondary metabolites in high quantities under controlled conditions (26,27) by reducing the formation of fungal agglomerates (28), thus leading to more efficient substrate consumption, greater oxygen transfer, and increased productivity of the fungus. Among Semi-SSF techniques, microparticle-enhanced cultivation (MPEC) with talcum powder or aluminum oxide (29) is widely used. When microparticles are added to culture media, some fungi tend to attach to them, inducing changes in the metabolism and as a result, the pattern of metabolites production can be modified (26,27,30,31).

In this work we set out to genomically characterize strain Aa22 in order to obtain insights into the genetic determinants of its ability to produce bioactive compounds, while obtaining a clear-cut taxonomic placement for the strain. We also tested fermentation conditions including solid (rice) and PDB liquid culture media without/with the addition of the microparticle-enhanced cultivation (MPEC) talcum as a potential inductor of metabolic changes. The ethyl acetate extracts obtained were analyzed for their chemical composition searching for targeted bioactive compounds previously described in this strain.

## Materials and Methods

### Biological material

The endophytic fungus Aa22 was isolated from *Artemisia absinthium* leaves. This strain was previously identified as *Stemphylium solani* and deposited in the Spanish Type Culture Collection (CECT 20941) (16).

### DNA extraction and size selection

High molecular weight fungal DNA was isolated using a modified cetyltrimethylammonium bromide (CTAB) method (32). Briefly, portions of approximately 150 mg of fungal mycelium grown in PDB medium for 3 days were washed with distilled water, centrifuged at 16,000 x g 1 min, and the pellet was ground to a fine powder in liquid nitrogen. The powder was mixed with 500 μL of CTAB buffer and 1.2 μL of 2-mercaptoethanol, followed by the addition of 2 μL of RNase A (10 mg/mL). After incubation at 65°C for 30 min with periodic mixing, proteins and lipids were removed by extraction using a chloroform-isoamyl alcohol mixture (24:1), with centrifugation at 17,300 x g for 10 min. The aqueous phase was transferred, and DNA was precipitated with 550 μL of cold isopropanol and 1/10 volume of 3M sodium acetate, followed by incubation at -20°C for 15 min. The DNA was pelleted by centrifugation at 9,500 x g for 10 min at 4°C, washed sequentially with 100% and 70% ethanol, air-dried and resuspended in EB buffer. DNA was purified using a size selection buffer to remove fragments under 10 kb (33), as follows. A 60 μL portion of DNA suspension was mixed with an equal volume of the size selection buffer. The mixture was centrifuged at 10,000 x g for 30 min at room temperature to pellet high-molecular-weight DNA, and the supernatant was carefully removed. The pellet was washed twice with 200 μL of 70% ethanol, centrifuging for 2 min at 10,000 g after each wash. The air-dried pellet was resuspended in EB buffer. DNA concentration and integrity were assessed using UV absorbance (Nanodrop) and fluorometry (Qubit). DNA purity was assessed based on 260/280 and 260/230 ratios, ensuring suitability for downstream applications.

### Genome sequencing, annotation and analysis

PacBio DNA HiFi libraries were prepared following the standard protocol and sequenced by Novogene (Cambridge, UK) on the Revio platform (PacBio, Menlo Park, CA). The genome was assembled and polished using Flye (v. 2.9); (34). Contigs were subjected to scaffolding with LRScaf (v. 1.1.10) with an identity threshold of 85%, a minimum overlap length of 500 bp, and a minimum overlap ratio of 60%. At least two supporting links were required, and repeat masking was applied. Genome sequences were annotated with Funannotate (v. 1.8.7; Palmer and Stajich 2020). Biosynthetic gene clusters (BGCs) were identified using the fungal version of antiSMASH (v. 8.0.1)(36). Taxonomic profiling was performed with UFCG (v. 1.0.6) (37), which extracts a predefined set of conserved fungal marker genes from the genome sequences. These markers were aligned and subsequently used to infer phylogenetic relationships. The phylogenetic tree was constructed with IQ-TREE (v2.3.1) (38) using the JTT (Jones– Taylor–Thornton) substitution model, incorporating empirical amino acid frequencies (F), a proportion of invariable sites (I), and a gamma distribution (G) to account for rate variation across sites. Branch support was evaluated through 1,000 bootstrap replicates. The resulting tree was visualized using FigTree (v1.4.4)(39)

Genome assembly quality was assessed with QUAST (v.5.2.0; (40), and completeness was evaluated with BUSCO (v.5.3.2) for *Ascomycota* and *Dothideomycetes* (41). Reference genomes and sequences used for identification were retrieved from publicly available data at NCBI (https://blast.ncbi.nlm.nih.gov/datasets/genome). Geneious (v. 9.1.8; (42) was used to build an Aa22 genomic database and to perform BLAST searches against our genome data. Protein sequences were aligned in NCBI (https://blast.ncbi.nlm.nih.gov/Blast.cgi). Functional domain annotation of protein sequences was conducted using InterProScan (https://www.ebi.ac.uk/interpro/search/sequence/).

### Fermentations

For solid-state fermentation (SSF) cultures were grown on rice grains. Firstly, inocula were grown on small Petri dishes (4 cm) with PDA medium (Difco, Maryland, USA). From these plates, sterile, 250-ml Erlenmeyer flasks containing 100 g of rice (commercial brand SOS Redondo, San Juan de Aznalfarache, Seville, Spain) and 50 ml of distilled water were inoculated with 6 mycelium fragments (1 cm^2^ each). Flasks were incubated at 25 °C in darkness for 7, 14, or 20 days (four replicates per time point).

For liquid-state fermentation PDB (BD-Difco, Sparks, MD) medium, with or without a MPEC physical support (talcum powder) was used. Cultures started from PDA plates. Sterile distilled water (10 mL) was added to each fully-grown PDA plate, its surface scraped with a spatula to obtain a mycelium suspension, and 2.5 mL of the suspension transferred to 4 500-mL Erlenmeyer flasks containing 100 mL PDB medium. Flasks were incubated in a rotary incubator (120 rpm; 25°C) in darkness, for 3 days. 20-mL portions of this preinoculum were used to inoculate 4 x 500-mL Erlenmeyer flasks containing 300 mL of PDB with or without talcum powder, and flasks were incubated under the same conditions for 7, 14, or 20 days (two replicates per time point). For MPEC cultures, talcum powder (Fisher Chemical, particle size 45 μm) was added at 6.6 g-L^-1^.

### Extract preparation

Mycelia were separated from culture media by paper filtration using a Büchner funnel. For SSF, filtered media were Soxhlet extracted (hexane followed by ethyl acetate). For liquid fermentations, sequential liquid-liquid extractions with hexane (to remove lipids) and ethyl acetate were carried out. After extraction, anhydrous Na_2_SO_4_ was added to the organic phase to remove any water traces, then filtered and rotary-evaporated at a reduced pressure.

Yields for liquid-liquid extractions were calculated as follows:

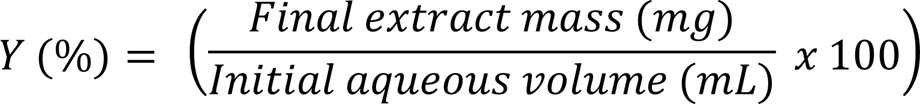

Where the final extract mass is the weight of the extract after solvent evaporation, and the initial aqueous volume is the amount of culture medium used for extraction.

### Gas Chromatography-Mass Spectrometry (GC-MS)

Analyses were conducted using a Shimadzu GC-2010 gas chromatograph equipped with a 30 m × 0.25 mm i.d. capillary column (0.25 μm film thickness, 95% Dimetil-5% diphenylpolisiloxane, Teknokroma TRB-5) coupled to a Shimadzu GCMS-QP2010 Ultra mass detector (electron ionization, 70 eV). Samples (1 μl) were injected with a Shimadzu AOC-20i autosampler. Working conditions were as follows: split ratio (20:1), injector temperature 300°C, temperature of the transfer line connected to the mass spectrometer 250 °C, initial column temperature 110 °C, then heated to 290 °C at 7 °C/min, and a full scan was used (m/z 35-450). Electron ionization mass spectra and retention data were used to assess the identity of compounds by comparing them with those found in the Wiley 229 and NIST 17 Mass Spectral Database and pure compounds isolated from Aa22 (16). All extracts (4 μg/μl) were dissolved in 100% dichloromethane for injection. Blank ethyl acetate extracts of all media were analyzed as controls.

## Results

### Genome sequence

Long genome sequence reads were obtained from high quality Aa22 DNA. A total of 5,160 Mb of HiFi sequence data was generated and assembled into 24 contigs, with a total genome size of 34.1 Mb and a mean coverage of 145x (S1 Table). The average GC content of 50.8%, and N50 and N90 values for this assembly were 1.79 Mb and 1.13 Mb respectively, indicating a high-quality assembly. Attempts to join contigs through scaffolding algorithms (see Materials and Methods) did not improve assembly (data not shown). The near-completeness of this genome sequence was confirmed by BUSCO analysis. For phylum Ascomycota, 1,647 complete BUSCOs were found out of 1,706 (96.5%), of which 1,644 were single-copy and 3 were duplicated. Twelve more were fragmented, and 47 (2.8%) were missing. For class Dothideomycetes, 3,625 BUSCOs were detected out of 3,786 (95.8%), with 3,621 as single-copy, 4 duplicated, 30 fragmented, and 131 (3.5%) missing. This genome has been deposited in NCBI under BioSample accession number SAMN47600574.

Availability of the near-complete Aa22 genome sequence allowed us to reappraise its taxonomic placement. Strain Aa22 had been classified as *Stemphylium solani* on the basis of the identity of a 580 bp sequence containing an incomplete ITS1-5.8S rRNA-ITS2 region with that of a *S. solani* strain (16). A BLASTN search of this sequence against the core-nucleotide BLAST database revealed 46 sequences with 100% identity (data not shown). Only two of them were *S. solani* (out of 126 *S. solani* sequences in the database), 41 were *S. lycopersici* (out of 201 *S. lycopersici* sequences in the database), and 3 were unclassified (data not shown). This raised the possibility that the two *S. solani* strains whose sequences were 100% identical to that of Aa22 may have been misclassified and that Aa22 was indeed a *S. lycopersici* strain. The Aa22 genome data, together with those of all available *Stemphylium* genomes at NCBI, were used for phylogenomic reconstruction (Fig. 1). Strain Aa22 clustered with three *S. lycopersici* genomes (strains CIDEFI212, CIDEFI213, and CIDEFI216) in a strongly supported clade (bootstrap value= 100), indicating close evolutionary relatedness to this species. Within this clade, Aa22 was more closely related *to S. lycopersici* CIDEFI212, with relatively high consistency (bootstrap value = 73). This clade was clearly distinct from other *Stemphylium* species for which genome sequences are available, such as *S*. *vesicarium* and *S. beticola*, which formed separate groups within the phylogeny (Fig. 1). However, no *S. solani* genomes have been sequenced to date. Taken together, the available evidence did not allow assigning strain Aa22 to either *S. solani* or *S. lycopersici*, two phytopathogenic species that appear to be closely related.

**Fig. 1.**
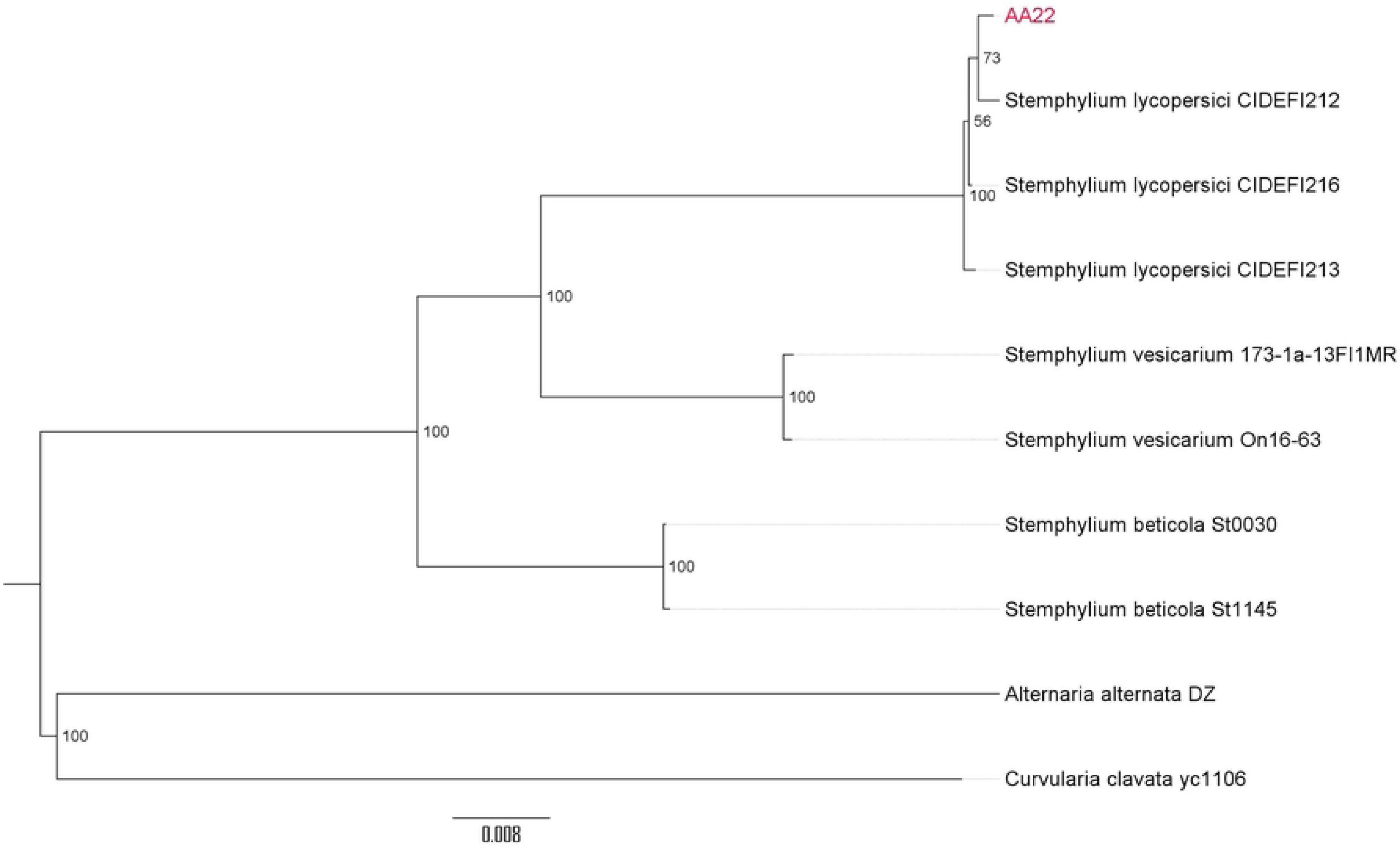
Phylogenomic tree of *Stemphylium* species. *Alternaria alternata* DZ and *Curvularia clavata* yc1106 were used as outgroups. Genomes were retrieved from the NCBI database (S2 Table). Figures indicate the % bootstrap support (1,000 replicates) for each node in the tree.

### Biosynthetic potential of the Aa22 genome

Analysis of the Aa22 genomic sequence with antiSMASH Fungi (https://fungismash.secondarymetabolites.org/) identified 42 gene coding regions associated with the production of secondary metabolites (Fig. 2, S3 table). Among these, one-third (14) of the regions displayed a varying degree of similarity to BGCs identified in other fungi, including 7 with low similarity, 3 with medium similarity and 4 with high similarity. In contrast, the remaining two-thirds (28) of the regions showed no clear match to known biosynthetic pathways. A total of 16 regions were linked to terpene biosynthesis, while 8 regions corresponded to polyketide synthesis (type I and type III polyketide synthases), and 7 were predicted to produce non-ribosomal peptides (NRPs). Additionally, 5 regions were associated with RiPP-like pathways (ribosomally synthesized and post-translationally modified peptides). The remaining six regions represented less common biosynthetic types, such as isocyanides and indole derivatives, with one region predicted for each. Seventeen of the regions contained hybrid BGCs, associated with more than one type of compound. Specifically, three of them involved terpene biosynthesis, seven contained polyketide synthases, four encoded non-ribosomal peptide synthases, and three contained RiPP-like regions.

**Fig. 2.**
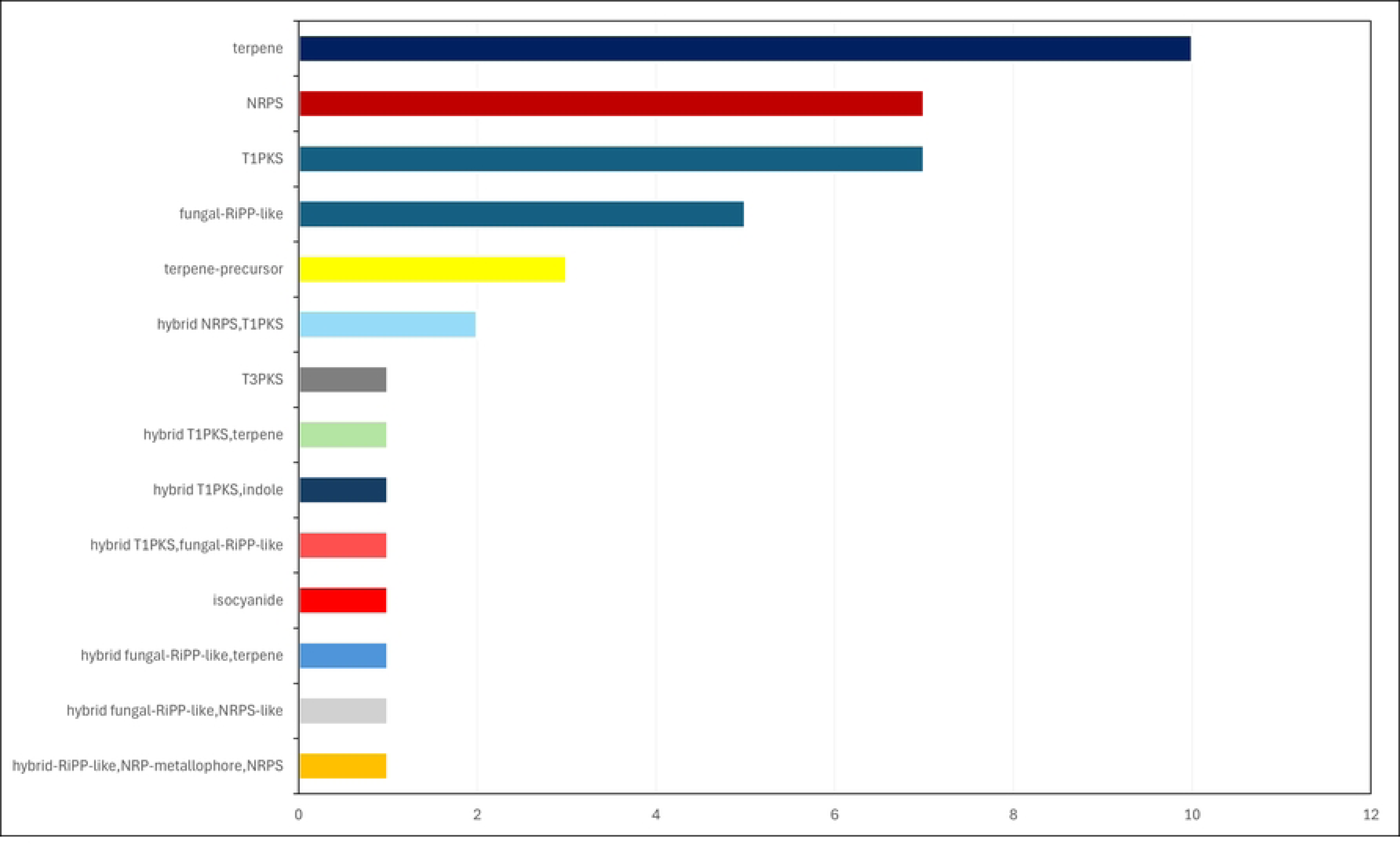
BGCs associated with *Stemphylium* sp. isolate Aa22. Their location and specific characteristics are shown in S3 Table.

The bio-active compounds produced by strain Aa22 (stemphol and stempholones A and B; (16) are alkyl resorcinols. Type III polyketide synthases (T3PKSs) have been implicated in the biosynthesis of alkyl-resorcinols (43–45). These enzymes use coenzyme A-bound substrates to form polyketide chains of varying length. They catalyze, within the same active site, the chain initiation, chain elongation, and cyclization reactions. Three different cyclization mechanisms have been described, giving rise to pyrone-quinolone- or resorcinol-polyketides (46,47). The sole T3PKS identified by antiSMASH Fungi in the Aa22 genome (Fig. 2) was further analyzed. A T3PKS from an endophytic *Fusarium incarnatum* isolate (GenBank accession KY780629.1), involved in the biosynthesis of resorcinol-like compounds (48), was used to BLAST search for similar regions in the Aa22 genome. A single significant match was identified within contig 16, where antiSMASH had predicted a T3PKS gene (S1 Fig.).

Pairwise sequence alignment showed 31.1% identity and 59.0% similarity between the *Fusarium* and *Stemphylium* proteins (S1 File). In addition, functional annotations using InterProScan confirmed both *Fusarium* and *Stemphylium* proteins within the T3PKS family, identifying N-terminal (IPR001099) and C-terminal (IPR012328) chalcone/stilbene synthase domains (S3 Table).

### Production of bio-active compounds

Fermentation yields in PDB are shown in Fig. 3A. In liquid fermentation, the maximum yields were obtained in the absence of talcum powder. The solid fermentation showed a yield increase with time (2.85, 180.0 and 252.4 mg/100g after 7, 14 and 20 days, respectively).

**Fig. 3.**
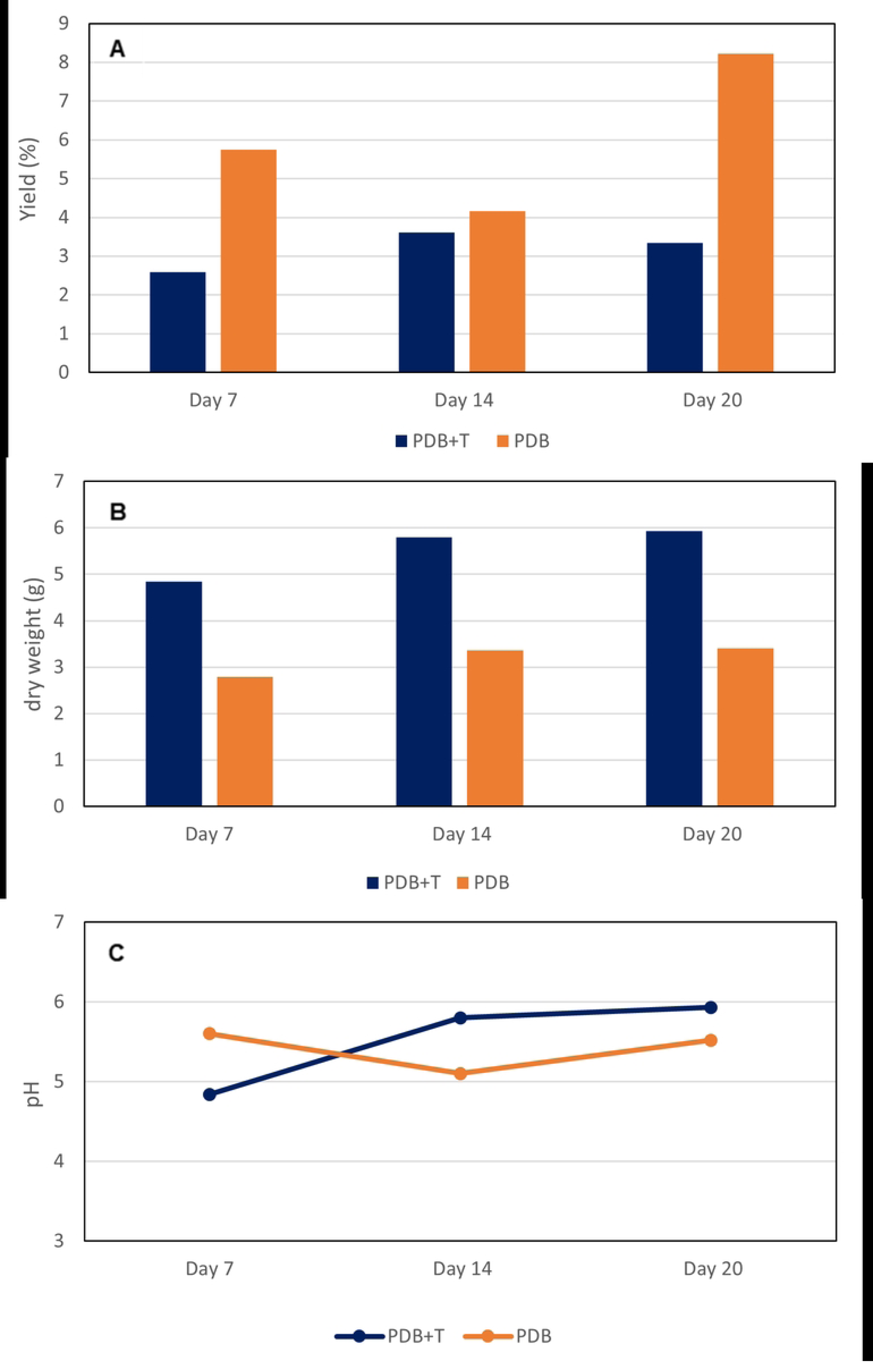
Submerged fermentation of strain Aa22 in PDB medium, with or without the addition of talcum powder. A: Solid extract yield; B: mycelium dry weight per fermentation; C: pH of the medium.

Higher mycelium growth was observed in fermentations with PDB (Fig. 3B). The pH (Fig. 3C) remained slightly acidic with minimal variation from the initial pH of 5.1, averaging *ca.* 6. In the case of fermentations with talcum powder, the pH was slightly higher (initial 5.77) probably due to the magnesium ions alkalizing the medium.

PDB cultures showed a characteristic, progressive darkening over time in PDB without talcum (Fig. 4).

**Fig. 4.**
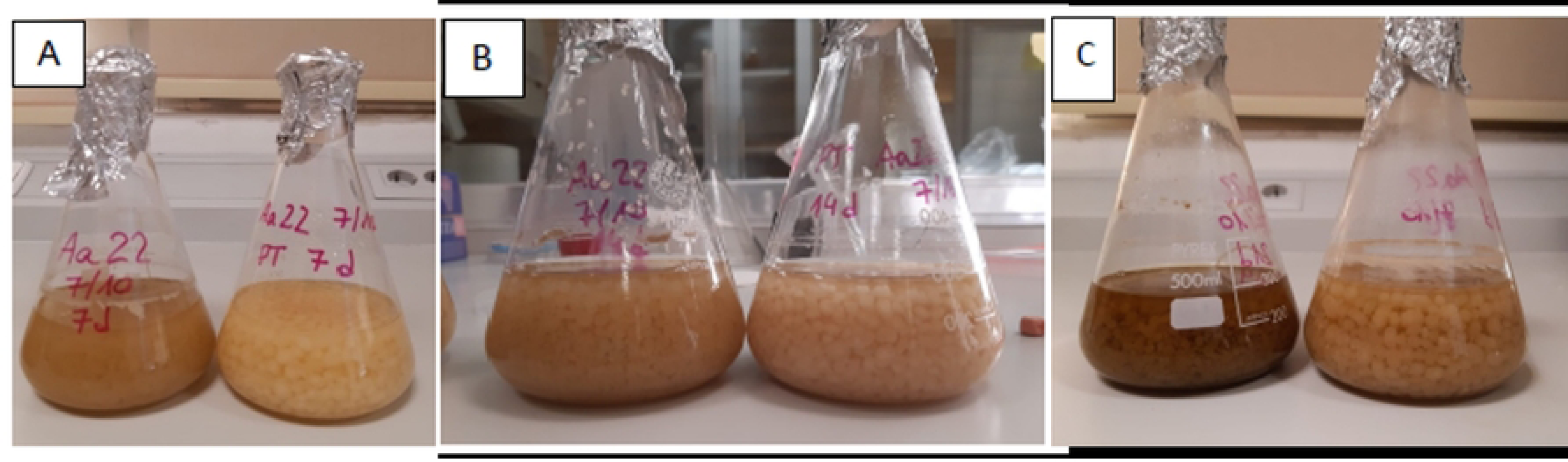
*Stemphylium* sp. Aa22 growth in fermentation flasks. PDB (left) and PDB plus talcum (right) after 7 (A), 14 (B), and 21 (c) days.

### Chemical Characterization

GC-MS analyses of extracts are presented in Table 3. Rice medium extracts were rich in fatty acids and derivatives, and their relative abundances changed with fermentation time. Major peaks were oleamide (10-18% with a peak at 14d), methyllinoleate (5-12% with a peak at 7d) and 9,12-hexadecadienoic acid, methyl ester (12-0.7% with a peak at 7d). No alkyl-resorcinols were detected. A mock rice extract had palmitic acid (26%), methyllinoleate (30%) and 8-heptadecenoic acid (28%) as the major components (S4 Table), suggesting that the lipid-rich composition of the solid-state medium extracts derives from rice lipids and their metabolism.

**Table 3.** GC-MS analysis of the Aa22 extracts fermented in Rice, PDB or PDB plus talcum media (7, 14 and 20 days). Data are shown as percent abundance.

| Retention time (min) | Compound | Rice |  |  | PDB |  |  | PDB+T |  |
| --- | --- | --- | --- | --- | --- | --- | --- | --- | --- |
|  |  | 7 d | 14 d | 20 d | 7 d | 14 d | 20 d | 7 d | 14 d |
| 3.51 | Phenylethyl alcohol |  |  |  | 1.35 | 3.92 |  | 11.02 |  |
| 4.22 | 4-(1-hydroxy-ethyl) γ-butanolactone | | | | 4.69 | | | 3.69 | |
| 7.34 | 1,2-Ethanediol |  |  |  | 3.68 | 4.57 |  | 5.02 |  |
| 7.66 | N-Butylacetamide |  |  |  | 5.13 |  |  |  |  |
| 11.39 | Acrylic acid tetradecanyl ester | 1.10 | 2.79 | 1.33 | 1.35 | 1.12 | 0.41 | 5.06 | 6.73 |
| 14.95 | Methyl palmitate | 4.56 | 2.84 | 0.80 | 1.79 |  |  |  |  |
| 17.41 | 9,12-Hexadecadienoic acid, methyl ester | 12.09 | 2.24 | 0.76 |  |  |  |  |  |
| 17.50 | Methyloleate | 9.10 | 1.97 | 0.97 |  |  |  |  |  |
| 17.94 | Methylinoleate | 12.37 | 7.23 | 5.20 |  |  |  |  |  |
| 18.58 | Palmitamide | 1.22 | 3.26 | 1.65 |  |  |  |  |  |
| 20.88 | Oleamide | 9.87 | 18.82 | 12.69 | 1.73 | 1.85 |  | 13.91 | 17.94 |
| 20.96 | 275/43/179/139/109/95/135/137/55/123 |  |  |  | 13.95 | 14.65 | 28.07 |  |  |
| 21.04 | Stempholone A |  |  |  | 9.56 | 11.30 | 22.52 |  |  |
| 21.44 | 3α-Hydroxy-manool | | | | 2.54 | | 4.25 | | |
| 24.37 | 190/43/107/137/95/245/81/121/55/109 |  |  |  | 8.16 | 20.88 | 16.76 |  |  |
| 24.7/29.6 | n-Hexatriacontane |  |  |  | 0.97 |  |  | 8.15 | 3.27 |
| 26.80 | Viridiflorol |  |  |  | 3.61 |  | 6.57 |  |  |
| 29.79 | 1-Eicosanol |  |  |  |  |  |  |  | 5.07 |

PDB extracts contained, especially at early times, low molecular weight volatiles (<200 g.mol^-1^) such as phenylethyl alcohol (0-4%, peak at 14d), 4-(1-hydroxyethyl)-butanolactone (0-4.7%, at 7d), 1.2-ethanediol (0-4.6%, at 7d) and N-butylacetamide (0-5.1%, at 7d) (Table 3). Two larger, unknown compounds were present at retention times of 20.96 min (28-14% with a peak at 21d) and 24.37 min (8-21% with a peak at 14d), together with stempholone A (9.5-22.5% with a peak at 20d). The presence of talcum powder (PDB+T) altered extract composition, with higher levels of phenylethyl alcohol at 7 days (11%) and accumulation of oleamide (14-18% with a peak at 7d) and other long-chain lipids. Probably as a result, the PDB+T extract at 21 days could not be analyzed due to its low solubility (Table 3). No detectable levels of stempholone A were found in PDB+T extracts.

Fig. 5 shows the composition-based dendrogram grouping of extracts. Two main groups were observed, matching the fermentation methods (solid and liquid).

**Fig. 5.**
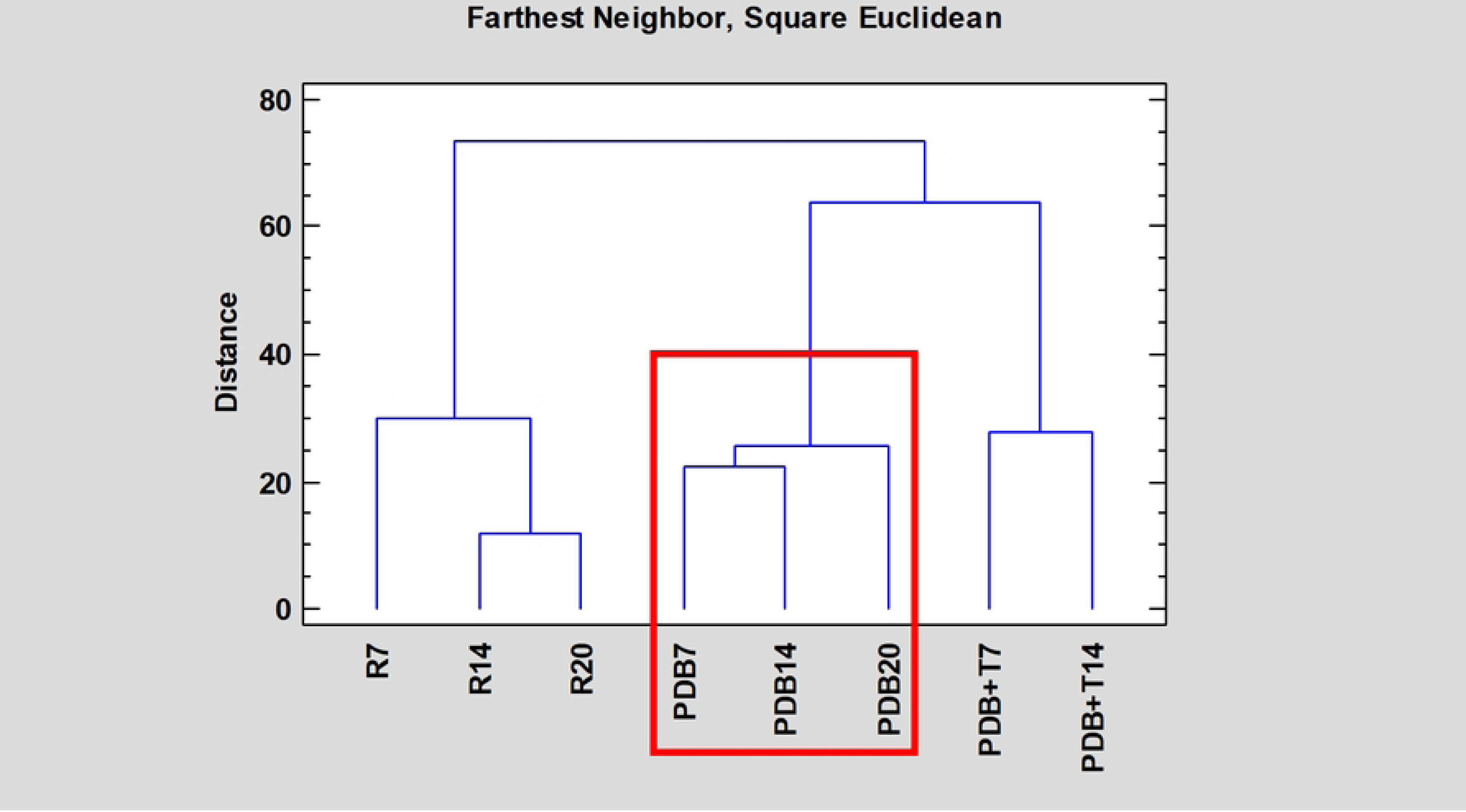
Dendrogram grouping of Aa22 extracts based on their chemical composition. Extracts containing Stempholone A are framed in red.

Additionally, two subgroups are observed for the liquid media (PDB and PDB+T). The PDB extracts were the only ones producing previously described bioactive compounds (stempholone A).

## Discussion

Whole genome sequencing and taxonomic profiling confirmed that isolate Aa22 represents a strain of the species *Stemphylium*, very close to *S. lycopersici*. However, given that some *S. solani* and *S. lycopersici* strains exhibit 100% identity in the ITS sequences originally used to taxonomically place strain Aa22 (16) and that no *S. solani* genome sequences are available, adscription of Aa22 to either species must remain uncertain. Both species are similar, and misclassification of strains has been reported as common (49). *S. lycopersici* and *S. solani* are common plant pathogens usually isolated from diseased plants. Strain Aa22 was isolated as an endophyte from a healthy wormwood plant (16). It is therefore noteworthy that its genome sequence is highly similar to those of pathogenic *S. lycopersici* isolates (Fig. 1), suggesting that minor differences exist at the genomic level between pathogenic and endophytic strains. In fact, besides strain Aa22, several endophytic isolates of both *Stemphylium* species have been isolated (50–53) suggesting that the endophytic lifestyle could be common among members of these species.

Aa22 has been previously reported as the producer of bioactive stempholones A and B and stemphol (antifeedants against the aphid *Myzus persicae*) (16). Stempholones A and B are produced in the same conditions as stemphol (16) and their biosynthetic pathway could be similar. Fungal biosynthetic pathways are complex, both in structure and in regulation (54). This complexity makes it difficult to associate specific genes or gene clusters with the production of certain metabolites during genome annotation (55). Regarding alkyl resorcinols, few T3PKSs specifically involved in their synthesis have been characterized (48). The identified Aa22 T3PKS shared the conserved domains identified in the *F. incarnatum* enzyme that was experimentally shown to synthesize alkyl resorcinols (48), suggesting a related function in polyketide biosynthesis. However, with only 31.1% sequence identity and 46.0% similarity, structural and functional differences are expected between these two enzymes, which could result in the production of different related metabolites (56). Given that only one T3PKS coding region was identified in the Aa22 genome, it is likely that its enzyme product plays a key role in the biosynthesis of stemphol, and possibly stempholones A and B. Future transcriptomic and mutagenesis studies will help clarify this role. Additionally, elucidating its regulatory mechanisms will provide insights into pathway activation, potentially enabling targeted induction strategies to enhance metabolite production as has been done in other studies (57).

The production of bio-active compounds by Aa22 cultures appeared to be linked to culture conditions. In this study, solid-state fermentation on rice produced extracts containing none of the described bioactive compounds extracts and rich in lipids derived from the rice substrate. Liquid fermentation in PDB medium resulted in higher extract yields with production of stempholone A along with two unidentified metabolites that increased with time. Addition of talcum powder (particle enhanced fermentation) decreased mycelium weight yield, an effect that had been observed in other filamentous fungi (58). For Aa22, fermentation yields also decreased with the addition of talcum powder. Talcum is a good adsorbent of molecules, with applications as a clarifier (59). This property may explain reduced extract yields. A different effect has been reported for an endophytic strain of *Phyllosticta capitalensis* (60) where talcum powder enhanced the production of meroterpenes, such as guignardone A and guignarenone B, at specific time points during fermentation, while the overall production of bioactive dioxolanones did not show significant improvements compared to the control. The effects of talcum powder on fermentation have also been linked to particle size. For instance, microparticles with a diameter of 40 μm improved the production of 2-phenyletanol by a strain of *Aspergillus terreus*, whereas smaller particles (10 μm) and iron oxide improved production of 2-phenyletanol and 6-pentyl-α-pyrone (58). Conversely, small talcum powder diameter (5 μm) resulted in the lowest polysaccharide yields among tested materials (61). These findings indicate that MPEC can significantly affect fermentations, either positively or negatively, across different organisms and metabolites. For Aa22, further research on MPEC is needed, either varying talcum powder diameter or using additional fermentation enhancers, such as bentonite or glass wool, as tested in other studies (60).

## Conclusions

The genome of *Stemphylium* sp. Aa22, an *Artemisia absinthium* endophytic strain previously described as a producer of the insect anti-feedant, alkyl-resorcinols stemphol and stempholones A and B, has been sequenced and analyzed. Genome-based taxonomic profiling placed strain Aa22 close to *S. lycopersici*. Forty-two BGCs, potentially responsible for biosynthesis of secondary metabolites, were identified. Among them, a sole type III polyketide synthase, potentially responsible for the biosynthesis of alkyl-resorcinols was identified.

Production of alkyl-resorcinols was strongly dependent on culture conditions. Solid fermentation on rice mainly produced lipid-based compounds, while fermentation in liquid PDB medium yielded two unidentified metabolites and the target compound stempholone A. In contrast, Micro-Particle Enhanced Cultivation (MPEC) adding talcum powder to PDB (PDB+T) resulted in a lower diversity of metabolites compared to PDB, with a higher proportion of lipid-derived compounds. Notably, previously reported bioactive compounds from this strain were not detected in solid or PDB+T fermentations. Further optimization of culture medium composition and fermentation conditions is required to maximize the biopesticidal potential of strain Aa22 and facilitate its scale-up.

## Author Contributions

**Conceptualization**: Azucena González-Coloma, Juan Imperial, Carmen Elisa Díaz, M. Fe Andrés.

**Data curation**: Jorge Rojas López-Menchero, Azucena González-Coloma, Juan Imperial, Carmen Elisa Díaz.

**Formal analysis**: Jorge Rojas López-Menchero, Azucena González-Coloma, Juan Imperial.

**Funding acquisition**: Azucena González-Coloma, Carmen Elisa Díaz.

**Investigation**: Jorge Rojas López-Menchero, Azucena González-Coloma, Juan Imperial, M. Fe Andrés.

**Methodology**: Jorge Rojas López-Menchero, Azucena González-Coloma, Juan Imperial, Carmen Elisa Díaz.

**Project administration**: Azucena González-Coloma, Carmen Elisa Díaz.

**Resources**: Azucena González-Coloma, Carmen Elisa Díaz, M. Fe Andrés.

**Supervision**: Azucena González-Coloma, Juan Imperial, Carmen Elisa Díaz.

**Validation**: Jorge Rojas López-Menchero, Azucena González-Coloma, Juan Imperial, Carmen Elisa Díaz.

**Visualization**: Jorge Rojas López-Menchero, Azucena González-Coloma.

**Writing – original draft**: Jorge Rojas López-Menchero, Azucena González-Coloma, Carmen Elisa Díaz.

**Writing – review & editing**: Jorge Rojas López-Menchero, Azucena González-Coloma, Juan Imperial, Carmen Elisa Díaz, M. Fe Andrés.

All authors have read and agreed to the published version of the manuscript.

## Acknowledgements

Access to computational resources was granted by the Galician Supercomputing Center (CESGA) through its supercomputing infrastructure. The supercomputer FinisTerrae III and its permanent data storage system have been funded by the Next Generation EU 2021 Plan de Recuperación, Transformación y Resiliencia, ICT2021-006904, and also from Programa Operativo Plurirregional de España 2014-2020 of the European Regional Development Fund (ERDF), ICTS-2019-02-CESGA-3, and from Programa Estatal de Fomento de la Investigación Científica y Técnica de Excelencia del Plan Estatal de Investigación Científica y Técnica y de Innovación 2013-2016 (1) Subprograma estatal de infraestructuras científicas y técnicas y equipamiento ERDF, CESG15-DE-3114

## Supporting information

**S1 Table. Sequencing and assembly statistics of the Stemphylium sp. isolate AA22 genome**

**S2 Table. Fungal genome assemblies retrieved from the NCBI database and used in this study**

**S3 Table. AntiSMASH Fungi prediction of regions containing Biosynthetic Gene Clusters (BGCs) for production of Secondary Metabolites**

**S4 Table. GC-MS analysis of mock extract obtained from rice.** Data are shown as percent abundance.

**S1 Fig. Gene organization of the region 8.1 in contig 16 predicted by antiSMASH containing the T3PKS coding region.**

**S1 file. Pairwise alignment of Fusarium incarnatum BMER1 T3PKS (GenBank accession KY780629.1) and the predicted T3PKS cluster (FUN_008199) from strain Aa22.**

## Notes

### Competing Interest Statement

The authors have declared no competing interest.

## References

1. Baker BP, Green TA, Loker AJ. Biological control and integrated pest management in organic and conventional systems. Biol Control. 2020 Jan;140:104095.

2. Kumar S, Singh A. Biopesticides: Present Status and the Future Prospects. J Fertil Pestic [Internet]. 2015 [cited 2025 Feb 4];06(02). Available from: https://www.omicsonline.org/open-access/biopesticides-present-status-and-the-future-prospects-jbfbp-1000e129.php?aid=64583

3. Damalas C, Koutroubas S. Current Status and Recent Developments in Biopesticide Use. Agriculture. 2018 Jan 12;8(1):13.

4. Samada LH, Tambunan USF. Biopesticides as Promising Alternatives to Chemical Pesticides: A Review of Their Current and Future Status. OnLine J Biol Sci. 2020 Feb 1;20(2):66–76.

5. Aly AH, Debbab A, Proksch P. Fungal endophytes: unique plant inhabitants with great promises. Appl Microbiol Biotechnol. 2011 Jun;90(6):1829–45.

6. Hu MY, Zhong GH, Sun ZhT, Sh G, Liu HM, Liu XQ. Insecticidal activities of secondary metabolites of endophytic *Pencillium* sp. in *Derris elliptica* Benth. J Appl Entomol. 2005 Sep;129(8):413–7.

7. Andrés MF, Diaz CE, Giménez C, Cabrera R, González-Coloma A. Endophytic fungi as novel sources of biopesticides: the Macaronesian Laurel forest, a case study. Phytochem Rev. 2017 Oct 24;16(5):1009–22.

8. Tawfike AF, Tate R, Abbott G, Young L, Viegelmann C, Schumacher M, et al. Metabolomic Tools to Assess the Chemistry and Bioactivity of Endophytic *Aspergillus* Strain. Chem Biodivers. 2017 Oct;14(10):e1700040.

9. Svobodová M, Šmídová K, Hvězdová M, Hofman J. Uptake kinetics of pesticides chlorpyrifos and tebuconazole in the earthworm Eisenia andrei in two different soils. Environ Pollut. 2018 May;236:257–64.

10. Strobel G, Daisy B, Castillo U, Harper J. Natural Products from Endophytic Microorganisms. J Nat Prod. 2004 Feb 1;67(2):257–68.

11. Morales-Sánchez V, Fe Andrés M, Díaz CE, González-Coloma A. Factors Affecting the Metabolite Productions in Endophytes: Biotechnological Approaches for Production of Metabolites. Curr Med Chem. 2020 Apr 23;27(11):1855–73.

12. Morales-Sánchez V, Díaz CE, Trujillo E, Olmeda SA, Valcarcel F, Muñoz R, et al. Bioactive Metabolites from the Endophytic Fungus Aspergillus sp. SPH2. J Fungi. 2021 Feb 2;7(2):109.

13. Bielecka M, Pencakowski B, Nicoletti R. Using Next-Generation Sequencing Technology to Explore Genetic Pathways in Endophytic Fungi in the Syntheses of Plant Bioactive Metabolites. Agriculture. 2022 Jan 28;12(2):187.

14. Chiquito-Contreras CJ, Meza-Menchaca T, Guzmán-López O, Vásquez EC, Ricaño-Rodríguez J. Molecular Insights into Plant–Microbe Interactions: A Comprehensive Review of Key Mechanisms. Front Biosci-Elite. 2024 Mar 12;16(1):9.

15. Batiha GES, Olatunde A, El-Mleeh A, Hetta HF, Al-Rejaie S, Alghamdi S, et al. Bioactive Compounds, Pharmacological Actions, and Pharmacokinetics of Wormwood (Artemisia absinthium). Antibiotics. 2020 Jun 23;9(6):353.

16. Diaz CE, Andres MF, Lacret R, Cabrera R, Gimenez C, Kaushik N, et al. Antifeedant, antifungal and nematicidal compounds from the endophyte Stemphylium solani isolated from wormwood. Sci Rep. 2024 Jun 12;14(1):13500.

17. Suriya J, Bharathiraja S, Krishnan M, Manivasagan P, Kim SK. Marine Microbial Amylases. In: Advances in Food and Nutrition Research [Internet]. Elsevier; 2016 [cited 2025 Feb 4]. p. 161–77. Available from: https://linkinghub.elsevier.com/retrieve/pii/S1043452616300341

18. Sharma A, Gupta V, Khan M, Balda S, Gupta N, Capalash N, et al. Flavonoid-rich agro-industrial residues for enhanced bacterial laccase production by submerged and solid-state fermentation. 3 Biotech. 2017 Jul;7(3):200.

19. Ouedraogo JP, Tsang A. Production of Native and Recombinant Enzymes by Fungi for Industrial Applications. In: Encyclopedia of Mycology [Internet]. Elsevier; 2021 [cited 2025 Feb 4]. p. 222–32. Available from: https://linkinghub.elsevier.com/retrieve/pii/B9780128199909000469

20. Sala A, Vittone S, Barrena R, Sánchez A, Artola A. Scanning agro-industrial wastes as substrates for fungal biopesticide production: Use of Beauveria bassiana and Trichoderma harzianum in solid-state fermentation. J Environ Manage. 2021 Oct;295:113113.

21. Ramos OS, Malcata FX. Food-Grade Enzymes. In: Comprehensive Biotechnology [Internet]. Elsevier; 2011 [cited 2025 Feb 4]. p. 555–69. Available from: https://linkinghub.elsevier.com/retrieve/pii/B9780080885049002130

22. Moresi M, Parente E. FERMENTATION (INDUSTRIAL) | Production of Some Organic Acids (Citric, Gluconic, Lactic, and Propionic). In: Encyclopedia of Food Microbiology [Internet]. Elsevier; 2014 [cited 2025 Feb 4]. p. 804–15. Available from: https://linkinghub.elsevier.com/retrieve/pii/B9780123847300001117

23. Seyed Reihani SF, Khosravi-Darani K. Mycoprotein Production from Date Waste Using Fusarium venenatum in a Submerged Culture. Appl Food Biotechnol [Internet]. 2019 Jan 22 [cited 2025 Feb 4];5(4). Available from: 10.22037/afb.v5i4.23139

24. Iram A, Özcan A, Yatmaz E, Turhan İ, Demirci A. Effect of Microparticles on Fungal Fermentation for Fermentation-Based Product Productions. Processes. 2022 Dec 13;10(12):2681.

25. Borah A, Selvaraj S, Murty VR. Production of Gallic Acid from Swietenia macrophylla Using Tannase from Bacillus Gottheilii M2S2 in Semi-Solid State Fermentation. Waste Biomass Valorization. 2023 Aug;14(8):2569–87.

26. Barrios-González J. Solid-state fermentation: Physiology of solid medium, its molecular basis and applications. Process Biochem. 2012 Feb;47(2):175–85.

27. Francis F, Druart F, Mavungu JDD, De Boevre M, De Saeger S, Delvigne F. Biofilm Mode of Cultivation Leads to an Improvement of the Entomotoxic Patterns of Two Aspergillus Species. Microorganisms. 2020 May 11;8(5):705.

28. Kowalska A, Boruta T, Bizukojć M. Morphological evolution of various fungal species in the presence and absence of aluminum oxide microparticles: Comparative and quantitative insights into microparticle-enhanced cultivation (MPEC). MicrobiologyOpen. 2018 Oct;7(5):e00603.

29. Walisko R, Krull R, Schrader J, Wittmann C. Microparticle based morphology engineering of filamentous microorganisms for industrial bio-production. Biotechnol Lett. 2012 Nov;34(11):1975–82.

30. Gibbs PA, Seviour RJ, Schmid F. Growth of Filamentous Fungi in Submerged Culture: Problems and Possible Solutions. Crit Rev Biotechnol. 2000 Jan;20(1):17– 48.

31. Bajoul Kakahi F, Ly S, Tarayre C, Deschaume O, Bartic C, Wagner P, et al. Modulation of fungal biofilm physiology and secondary product formation based on physico-chemical surface properties. Bioprocess Biosyst Eng. 2019 Dec;42(12):1935–46.

32. Doyle J. DNA Protocols for Plants. In: Hewitt GM, Johnston AWB, Young JPW, editors. Molecular Techniques in Taxonomy [Internet]. Berlin, Heidelberg: Springer Berlin Heidelberg; 1991 [cited 2025 Mar 31]. p. 283–93. Available from: http://link.springer.com/10.1007/978-3-642-83962-7_18

33. Jones A, Purushotham N, Nasim J, Schwessinger B. DNA clean-up and size selection for long-read sequencing v4 [Internet]. 2021 [cited 2025 Feb 4]. Available from: https://www.protocols.io/view/dna-clean-up-and-size-selection-for-long-read-sequ-bwkdpcs6

34. Kolmogorov M, Yuan J, Lin Y, Pevzner PA. Assembly of long, error-prone reads using repeat graphs. Nat Biotechnol. 2019 May;37(5):540–6.

35. Palmer JM, Stajich J. Funannotate v1.8.1: Eukaryotic genome annotation [Internet]. Zenodo; 2020 [cited 2025 Feb 4]. Available from: https://zenodo.org/record/1134477

36. Blin K, Shaw S, Steinke K, Villebro R, Ziemert N, Lee SY, et al. antiSMASH 5.0: updates to the secondary metabolite genome mining pipeline. Nucleic Acids Res. 2019 Jul 2;47(W1):W81–7.

37. Kim D, Gilchrist CLM, Chun J, Steinegger M. UFCG: database of universal fungal core genes and pipeline for genome-wide phylogenetic analysis of fungi. Nucleic Acids Res. 2023 Jan 6;51(D1):D777–84.

38. Minh BQ, Schmidt HA, Chernomor O, Schrempf D, Woodhams MD, Von Haeseler A, et al. IQ-TREE 2: New Models and Efficient Methods for Phylogenetic Inference in the Genomic Era. Teeling E, editor. Mol Biol Evol. 2020 May 1;37(5):1530–4.

39. Rambaut A. FigTree v1.4.4 [Software]. Institute of Evolutionary Biology, University of Edinburgh; 2018. Available from: http://tree.bio.ed.ac.uk/software/figtree/

40. Gurevich A, Saveliev V, Vyahhi N, Tesler G. QUAST: quality assessment tool for genome assemblies. Bioinformatics. 2013 Apr 15;29(8):1072–5.

41. Manni M, Berkeley MR, Seppey M, Zdobnov EM. BUSCO: Assessing Genomic Data Quality and Beyond. Curr Protoc. 2021 Dec;1(12):e323.

42. Kearse M, Moir R, Wilson A, Stones-Havas S, Cheung M, Sturrock S, et al. Geneious Basic: An integrated and extendable desktop software platform for the organization and analysis of sequence data. Bioinformatics. 2012 Jun 15;28(12):1647–9.

43. Miyanaga A, Horinouchi S. Enzymatic synthesis of bis-5-alkylresorcinols by resorcinol-producing type III polyketide synthases. J Antibiot (Tokyo). 2009 Jul;62(7):371–6.

44. Katsuyama Y, Ohnishi Y. Type III Polyketide Synthases in Microorganisms. In: Methods in Enzymology [Internet]. Elsevier; 2012 [cited 2025 Mar 31]. p. 359–77. Available from: https://linkinghub.elsevier.com/retrieve/pii/B9780123942906000173

45. Milke L, Kabuu M, Zschoche R, Gätgens J, Krumbach K, Carlstedt KL, et al. A type III polyketide synthase cluster in the phylum Planctomycetota is involved in alkylresorcinol biosynthesis. Appl Microbiol Biotechnol. 2024 Dec;108(1):239.

46. Hertweck C. The Biosynthetic Logic of Polyketide Diversity. Angew Chem Int Ed. 2009 Jun 15;48(26):4688–716.

47. Shimizu Y, Ogata H, Goto S. Type III Polyketide Synthases: Functional Classification and Phylogenomics. ChemBioChem. 2017 Jan 3;18(1):50–65.

48. Manoharan G, Sairam T, Thangamani R, Ramakrishnan D, K.Tiwari M, Lee JK, et al. Identification and characterization of type III polyketide synthase genes from culturable endophytes of ethnomedicinal plants. Enzyme Microb Technol. 2019 Dec;131:109396.

49. Nasehi A, Al-Sadi AM, Nasr Esfahani M, Ostovar T, Rezaie M, Atghia O, et al. Molecular re-identification of Stemphylium lycopersici and Stemphylium solani isolates deposited in NCBI GenBank and morphological characteristics of Malaysian isolates. Eur J Plant Pathol. 2019 Mar;153(3):965–74.

50. Ali R, Gul H, Rauf M, Arif M, Hamayun M, Husna, et al. Growth-Promoting Endophytic Fungus (Stemphylium lycopersici) Ameliorates Salt Stress Tolerance in Maize by Balancing Ionic and Metabolic Status. Front Plant Sci. 2022 Jul 11;13:890565.

51. Husna H, Hussain A, Shah M, Hamayun M, Iqbal A, Qadir M, et al. Stemphylium lycopersici and Stemphylium solani improved antioxidant system of soybean under chromate stress. Front Microbiol. 2022 Nov 3;13:1001847.

52. Zhong L, Niu B, Xiang D, Wu Q, Peng L, Zou L, et al. Endophytic fungi in buckwheat seeds: exploring links with flavonoid accumulation. Front Microbiol. 2024 Feb 20;15:1353763.

53. Li Y, Zhu G, Wang J, Yu J, Ye K, Xing C, et al. New Polyketide Congeners with Antibacterial Activities from an Endophytic Fungus Stemphylium globuliferum 17035 (China General Microbiological Culture Collection Center No. 40666). J Fungi. 2024 Oct 24;10(11):737.

54. Naranjo-Ortiz MA, Gabaldón T. Fungal evolution: cellular, genomic and metabolic complexity. Biol Rev. 2020 Oct;95(5):1198–232.

55. Mohanta TK, Al-Harrasi A. Fungal genomes: suffering with functional annotation errors. IMA Fungus. 2021 Dec;12(1):32.

56. Morris R, Black KA, Stollar EJ. Uncovering protein function: from classification to complexes. Essays Biochem. 2022 Aug 10;66(3):255–85.

57. Sun WW, Li CY, Chiang YM, Lin TS, Warren S, Chang FR, et al. Characterization of a silent azaphilone biosynthesis gene cluster in Aspergillus terreus NIH 2624. Fungal Genet Biol. 2022 May;160:103694.

58. Gonciarz J, Bizukojc M. Adding talc microparticles to *A spergillus terreus* ATCC 20542 preculture decreases fungal pellet size and improves lovastatin production. Eng Life Sci. 2014 Mar;14(2):190–200.

59. Grafia AL, Castillo LA, Barbosa SE. Use of talc as low-cost clarifier for wastewater. Water Sci Technol. 2014 Feb 1;69(3):640–6.

60. Reyes Castillo N, Rojas López-Menchero J, Pacheco Useche WA, Díaz CE, Andres MF, González-Coloma A. Advanced fermentation techniques enhance dioxolanone type biopesticide production from Phyllosticta capitalensis. Sci Rep. 2025 Mar 7;15(1):7989.

61. Fan JH, Xiong LQ, Huang W, Hong JQ, Guo HK, Wong KH, et al. Exopolysaccharides produced by Antrodia cinnamomea using microparticle-enhanced cultivation: Optimization, primary structure and antibacterial property. Int J Biol Macromol. 2024 Feb;259:128872.

